# Different responses to cell crowding determine the clonal fitness of *p53* and *Notch* inhibiting mutations in squamous epithelia

**DOI:** 10.1101/2020.03.30.015917

**Authors:** Vasiliki Kostiou, Michael WJ Hall, Philip H Jones, Benjamin A Hall

## Abstract

The growth and competition of cells in epithelial tissues plays an important role in both tissue homeostasis and the robustness of normal tissues to pre-cancer mutation. Whilst wild-type cells compete neutrally for dominance in the un-mutated tissue, naturally occurring mutations in individual cells may lend them a fitness advantage that can allow tissue colonisation. In mouse oesophageal epithelia, the growth of *p53* mutants and a dominant negative mutant of the *Notch* downstream target *Maml1* (*DN_Maml1*) have been shown to have different colonisation properties despite strong quantitative similarities in the growth of individual clones. Here we show that in order to recapitulate these behaviours whilst maintaining tissue turnover models need to take account of the response of cells to increased areal density in the tissue colonised by mutant cells. We demonstrate that *p53* mutant clone growth approximates a logistic curve, but that without including limitations on mutation induced expansion the overall proliferation rate of the tissue drops due to space restrictions. In contrast, the ability of *DN_Maml1* mutations to displace the wild-type population reflects a feedback that effects both mutant and wild-type cells equally. We go on to show how these distinct feedbacks are consistent with the distribution of mutations observed in human datasets.

## Introduction

Adult tissues rely on progenitor cells to maintain lifelong proper function of organs. A major subject of study in biological research is the dynamics of progenitor cells in squamous epithelia. These are rapidly regenerative tissues covering the external surface of the body, the mouth, and the oesophagus and organised in layers of keratinocytes. Progenitor cells communicate and interact with one another and their environment in a tightly coordinated manner to respond to tissue needs and ensure homeostasis.

Progenitor cell dynamics has been shown to be accurately described by a simple mathematical model, the Single Progenitor (SP) model [1–4]. This proposes that a single, equipotent progenitor cell population maintains the tissue. In mouse stratified squamous epithelia, proliferation is restricted in the basal layer where progenitor cells stochastically divide or differentiate through stratifying into the upper layers of the tissue before eventually being shed (Fig 1a). Division probabilities are balanced, allowing homeostasis to be achieved across the progenitor population (Eq 1).

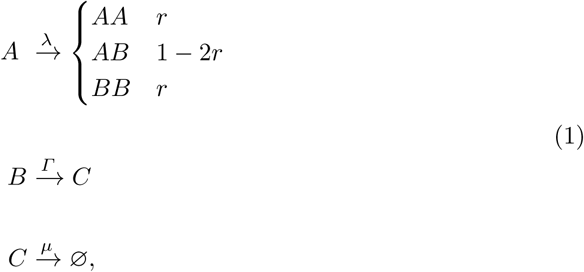

where *A* represents the basal layer progenitor cells, *B* the basal cells committed to differentiate and *C* the suprabasal layer cells. Progenitor cells divide regularly with an overall division rate λ and give rise to either two progenitor daughters (*AA*), two differentiating daughters (*BB*) or one daughter of each type (*AB*) with fixed probabilities. Given the fact that *AA* symmetric division leads to clone expansion and *BB* symmetric division tends towards to clone extinction, the two symmetric division rates should be equal in order for a steady state in terms of number of cells to be maintained across the progenitor clone population. The probabilities of symmetric and asymmetric divisions are *r* and 1 − 2*r* respectively with 0 < *r* < 0.5. Differentiating daughters in the basal layer stratify to the suprabasal layer at rate Γ and supra basal cells, *C*, are shed at rate *µ*.

**Fig 1.**
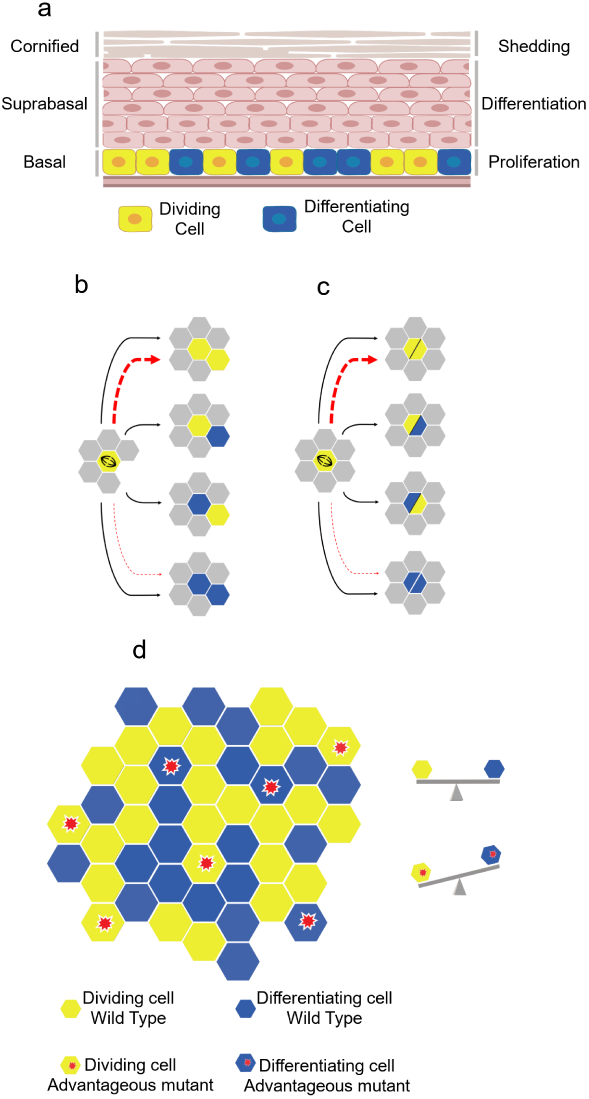
a: The architecture of murine stratified squamous epithelial tissues. Proliferation is restricted to the lowest basal layer. Upon differentiation, basal cells exit the cell cycle and migrate through suprabasal layers, until eventually, they are shed from the tissue. Cell production and loss should be perfectly balanced so that homeostasis and tissue proper function is achieved. Figure was generated using BioRender.com. b,c: Schematic representation of the rules of the spatial single progenitor model. Proliferating cells in yellow and differentiating cells in blue. Proliferating cells undergo a division type with balanced division outcome probabilities. In case of mutant cells, division probabilities are biased, favouring symmetric division, indicated by dashed red lines. A proliferating cell which is about to divide checks its immediate neighbourhood for available space. If a vacant site exists (b), one daughter cell occupies the mother cell’s space and the second the neighbouring empty space. If there is no empty space in the immediate neighbour (c), the two daughters occupy the mother cell’s space, thus creating a double cell occupancy. Double state cells are released once a neighbouring lattice site becomes available. d: Illustration of the two-dimensional hexagonal lattice representing the epithelial basal layer. Proliferating cells are able to divide, whilst differentiating cells exit the basal layer and are removed from the simulation. Mutant cells, marked with a red asterisk, have a bias in producing proliferating daughters.

In view of the fact that most common human cancers appear in epithelia [5], understanding the mechanisms of cell growth and competition within the spatial constraints of these tissues is fundamental to explain not only healthy tissue growth and maintenance but also the processes of mutagenesis and cancer. Within the continuous sheet of epithelium, cell populations constantly compete for space, minimizing redundant or insufficient cell production and sustaining homeostasis. This may be accomplished by mechanical interactions, e.g. through the mechanosensing ion channel Piezo1 [6, 7], and molecular signalling, e.g. through the *Notch* signalling pathway [8]. Whilst wild-type cells compete neutrally for space in the un-mutated tissue, naturally occurring mutations in individual cells may lend them a fitness advantage that can allow tissue takeover. Thus, non neutral competition between cells with different fitness levels leads to the dominance of the fitter population (“winners”) at the expense of the less fit population (“losers”). This is a mechanism by which oncogenic mutations colonize epithelial tissue and potentially drive preneoplasia and tumour formation.

Mutations in the tumour suppressor gene *p53* and *Notch* pathway have been shown to exhibit non neutral growth dynamics in epithelial tissues. *Maml1* is required in the canonical *Notch* pathway as it forms a complex with the *Notch* intracellular domain, thus enabling the transcription of target genes [22]. An increasing body of work suggests that *p53* mutations are highly frequent in squamous epithelial cancers [9–11]. Furthermore, recent studies report high incidence of *p53* and *Notch1* mutated genes in normal human skin and oesophagus [12–14]. This highlights the importance of studying the process of accumulation and interaction of these mutations within tissues in order to understand the mutational landscape from which cancer develops. Two recent studies investigating the growth dynamics of advantageous mutations in mouse oesophagus and back skin epidermis respectively, have shown that Notch pathway mutants fully colonize the tissue whereas *p53* mutants do so only partially. Here, using a computational model of clonal competition in the tissue, we demonstrate that the observed difference in mutant clone dynamics can be explained by different spatial feedback rules, on the basis of response to cell crowding. Several lines of evidence support the response of epithelial cells to crowding (i.e. increased areal density) as a spatial regulatory mechanism between surrounding cells. Previous studies in epithelial tissues of different model organisms argue that local crowding and cell deformation induced by proliferation could trigger the delamination of nearby non-apoptotic cells [7, 17, 18]. Furthermore, a recent study on mouse plantar epidermis suggests that division is stimulated by a stratification event in the local environment [19]. Our findings imply that the two mutant populations show a diverse response to crowding, providing a mechanistic explanation of the observed distinct growth modes.

## Materials and methods

### Spatial single progenitor model

A stochastic cellular automaton (CA) was used to implement the SP model in two-dimensional space and explore the collective behaviour of cells in the tissue. A two-dimensional, hexagonal lattice was used to model the basal layer of the epithelium, reflecting the observation that each oesophageal basal cell has six neighbours on average (Fig 1d). Basal epithelial cells were simulated on a lattice originally containing 10,000 cells (*L* = 100 *X* 100), corresponding to roughly 1% of the area of adult mouse oesophagus, where periodic boundary conditions were applied. Each simulation was repeated 100 times.

Each site of the grid may be occupied either by one of the two cell types described in the SP model, proliferating cells (A) and post-mitotic cells (B), or it may remain vacant as a result of a stratification event. Also, a lattice site may be occupied by two cells, indicating a crowded region. A division event can lead to three potential outcomes: two proliferating cells, two differentiating cells or one daughter of each cell type. The neighbourhood in the SP CA model is defined by the six adjacent places. Division and stratification events were considered as two independent processes determined solely by r, λ and Γ parameters. This cell-autonomous approach could lead to cases where a cell division event occurs at a region with no available neighbouring vacant space. In this case, the two daughter cells were placed on the same grid space, indicating an increased cell density area. Analogously, cases where an empty space generated by a recently stratified B cell is not rapidly replaced by a nearby newly born cell might be observed, representing a low cell density area. Considering the above, each lattice site could have one of the following seven potential states: *A, B, D*_*AA*_, *D*_*AB*_, *D*_*BA*_, *D*_*BB*_ and “empty” (∅), where *D*_*AA*_, *D*_*AB*_, *D*_*BA*_, *D*_*BB*_ correspond to double occupancies (Fig 1b,c). Thus, the SP model (Eq 1) was extended to explicitly include space as follows:

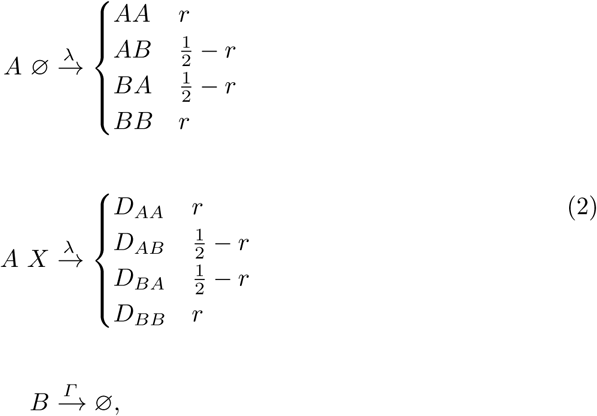

where *A* ∅ denotes a type A cell neighbouring a vacant lattice site and *AX* denotes a type A cell neighbouring either a type A or type B cell, thus indicating that there is no neighbouring empty space. *D*_*AA*_, *D*_*AB*_, *D*_*BA*_, *D*_*BB*_ correspond to double occupancies. The CA model was developed in NetLogo [15]. We used a Markovian stochastic simulation algorithm where the basal layer was simulated as an asynchronous CA. The algorithm included the following steps:

1. Start by defining a grid of *NxN* sites (*N* = 100) with A and B cells randomly seeded.
2. For every cell on each lattice site, draw a random number from an exponential distribution with mean 1/*λ* or 1/*Γ* to assign time of next event (division or stratification) for A and B cells respectively.
3. Select cell with the smallest next event time assigned. Current time is updated to the smallest next event time.
4. If an A cell is selected, use a random number from a uniform distribution *U* ∈ {0, 1} to choose the division type to occur by comparing *U* to division probabilities. Assign the division type as a next event for the selected cell. If a B cell is selected, assign stratification as a next event for the selected cell.
5. If the next event is division, all neighbouring places are checked for empty space. In the case of an existing neighbouring space, one new born cell will replace the mother cell and the other will occupy the empty neighbouring space. If there is no empty neighbouring space available then both will remain at the mother cell’s space (creating a “double state” cell), until a neighbouring space is released. If stratification is the next event, B cell stratifies, leaving an empty space, which allows potential neighbouring “double state” daughters to be released.
6. Repeat steps 3-6 until there are no A or B cells left or time threshold is reached.

### Spatial single progenitor model of non-neutral growth

The study of *p53* and *DN_Maml1* mutant clone dynamics in mouse epithelia indicated that the mutant progenitor clones are not in homeostasis in a mixed tissue, and have a fitness advantage over their wild-type counterparts [9, 16]. This has been shown to be achieved by having a bias towards the production of proliferating progeny. Such bias results in a gradual expansion in the proliferating population over time as there are less chances for the mutant clones to be lost by differentiation. Thus, mutant clonal expansion appears to be consistent with a SP model including a cell fate imbalance:

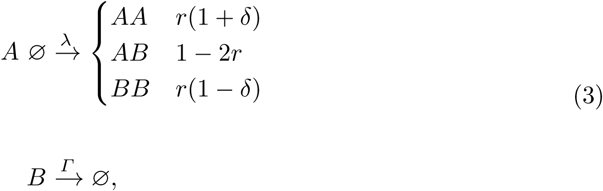

where *δ* denotes the tilt in cell fate. Therefore, *δ* = 0 corresponds to homeostasis, *δ* = 1 implies absence of symmetric differentiation, leading to persistence and *δ* = −1 implies absence of symmetric division, leading to extrusion.

To simulate mutant clonal dynamics in two-dimensional space, we used the spatial SP model including a fate imbalance, considering both wild-type and mutant epithelial cells. The choice of the grid architecture, the neighbourhood, the number of states and spatial rules of the CA model were implemented as described in the homeostatic spatial SP model. However, a new mutation-status property was introduced to distinguish mutant cells to wild-type cells. In the case of *p53* cells, mutants have the exact same properties as the wild-type cells and the only thing that distinguishes the two cell populations is that *p53* mutant cells have an innate bias towards the production of proliferating cells (*δ*) (Fig 1b,c). In the case of *DN_Maml1* mutant cells, mutants will also have distinct λ, Γ and *r* values, as inferred by [9]. Considering the above, the spatial SP model, initially described in Eq 2 was modified as follows, in order to accommodate the mutant cell population:

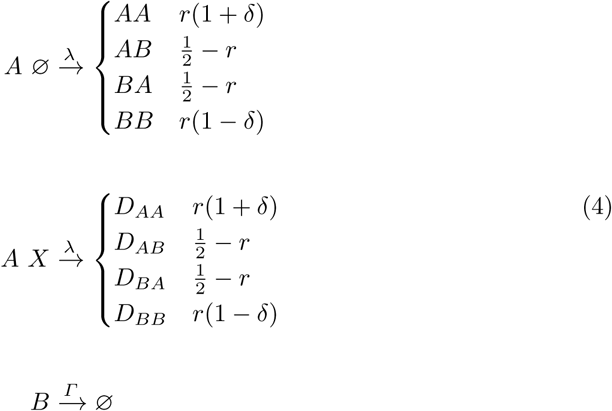

The simulation steps were modified as follows:

1. Start by defining a grid of *NxN* sites (*N* = 100) with A and B cells randomly seeded.
2. Insert randomly mutant cells of type A.
3. For every cell on each lattice site, draw a random number from an exponential distribution with mean 1/*λ* or 1/*Γ* to assign time of next event (division or stratification) for A and B cells respectively.
4. Select cell with the smallest next event time assigned. Current time is updated to the smallest next event time.
5. If an A cell is selected, use a random number from a uniform distribution *U* ∈ {0, 1} to choose the division type to occur by comparing *U* to division probabilities. Symmetric division probabilities for mutant cells are biased according to *δ*. Assign the division type as a next event for the selected cell. If a B cell is selected, assign stratification as a next event for the selected cell.
6. If the next event is division, all neighbouring spaces are checked for empty space. In the case of an existing neighbouring space, one new daughter cell will replace the mother cell and the other will occupy the empty neighbouring space. If there is no empty neighbouring space available then both will remain at the mother cell’s space (creating a “double state” cell), until a neighbouring space is released. If stratification is the next event, B cell stratifies, leaving an empty space, which allows a pair of potential neighbouring “double state” daughters to be released.
7. Repeat steps 4-7 until there are no A or B cells left or time threshold is reached.

To introduce feedbacks in the non-neutral simulations, we followed the same steps and modified the division probabilities. To perform simulations with the upstream feedback an additional fate bias parameter, *δ′*, was introduced. The value of *δ′* depends on the local cell density, i.e. number of neighbouring cells. A neighbourhood (*n*) consisting of more than a defined number of cells would be considered as “crowded” whereas fewer neighbouring cells than the defined crowding threshold would indicate an underpopulated region (”empty”). In the former case, *δ′* is negative rendering the fate of dividing cells tilted towards symmetric differentiation to release crowding. In the latter case, a positive *δ′* is chosen, favouring symmetric division to fill the empty sites. When the neighbourhood is neither “crowded” nor “empty”, this indicates that the local cell density is homeostatic and therefore *δ′* = 0.

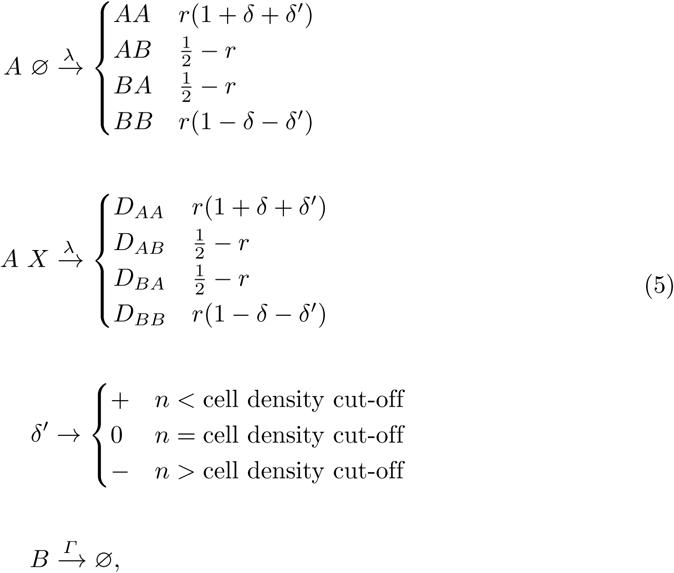

As each cell on the grid has six neighbours in normal conditions, i.e. no overcrowding, no gaps, the local cell density crowding cut-off was set to six cells. This reflects the topology of the healthy tissue. Furthermore, as wild type cells do not have a *δ*, they would solely respond to the cell density bias (*δ′*), whereas in the case of mutant cells the cell density bias (*δ′*) and their innate fate bias (*δ*) would be counterbalanced (Eq 5). To avoid large increases in overall tissue cell density, an additional rule was applied to every cell in the grid that had an overcrowded neighbourhood (*n* > 8 cells). In such case the probability of that cell to undergo symmetric division was minimized (*AA* = 0, *BB* = 2*r*).

To perform simulations with the downstream feedback mechanism, no additional cell density fate bias parameter (*δ′*) was used but mutants were set to lose their innate bias towards the production of proliferating progeny, (*δ*), in response to crowding in their local neighbourhood. Thus, the downstream feedback model can be described by the Eq 4 relationship of a spatial model with fate bias. The value of *δ* parameter is turned off when the local cell density (i.e. number of cells) of a dividing mutant cell’s neighbourhood consists of more than a defined crowding threshold, switching the mutant behaviour to balanced mode (*δ* = 0). Moreover, to allow for stress release induced by the double state cells observed in downstream feedback simulations, we introduced an additional rule where every new division that gave rise to a double occupancy consisting of at least one differentiating cell (AB, BA or BB double state cells) would lead to an instantaneous stratification event, that is, the removal of one B cell from the simulation. This additional rule is consistent with previous reports of cell extrusion events to compensate proliferation induced local stress [7, 17, 18].

To perform mutant competition simulations both downstream and upstream feedback rules were introduced in the same simulation. *p53* mutant cells followed the downstream feedback whilst *DN_Maml1* mutants and wild-type cells followed the upstream feedback.

## Results

### Tissue organisation does not alter the neutral single progenitor model

Spatial competition plays an important role in a wide variety of different scenarios, altering growth curves in ways that reflect both tissue structure and experimental methods [20, 21]. The finite size of the basal layer of the tissue constrains clone growth, and the topology of the cells restrains clonal expansion to the periphery of the clone. These features potentially alter growth patterns and undermining the validity of the model. In the single progenitor model specifically we might expect that average clone size grows initially linearly, but becomes sublinear as clones grow large and only can expand through competition with neighbouring clones at the periphery. We therefore sought initially to confirm that the single progenitor model continues to match experimental data when spatial competition is explicitly included. The spatial single progenitor model was initially developed without accounting for any kind of spatial feedbacks that could influence cell fate decisions, similar to the non-spatial model.

The simulation outputs revealed that taking into consideration the spatial constraints imposed by the lattice do not cause the model to deviate from the experimental data taken from mouse oesophagus (Fig 2). The spatial model is able to reproduce the characteristic features of the stochastic birth-death process (the hallmarks of a single population of progenitor cells), as described formally in [3]. Homeostasis (a constant number of proliferating cells) is maintained by a balance between cell production and loss (Fig 2a). Over time, due to stochastic divisions resulting in a pair of differentiated daughter cells, clones are lost following a simple relationship determined by r and λ. To maintain a constant population, the average size of the persisting clones rises linearly (with slope *rλ*/*ρ*). We observe that the simulated tissue demonstrates both of these properties (Fig 2b,c) with growth dictated as expected by input model parameters. Finally, whilst the clone size distribution becomes broader over time, the shape remains constant once scaled by the average clone size (Fig 2d,e). Taken together, these findings demonstrate that the average behaviour and the clone size distributions of the epithelial basal cells are consistent with the SP model of epithelial homeostasis. We further extended this testing to all available datasets, and find that the growth patterns of the new model accurately reproduce experimental observations (S1 Fig).

**Fig 2.**
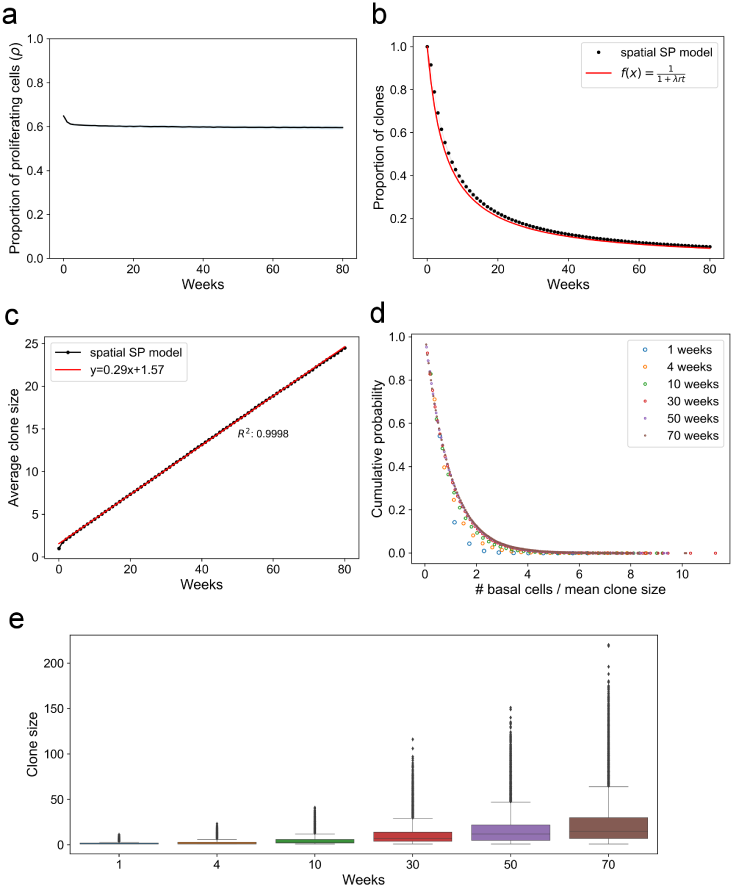
The single progenitor hallmarks on cell populations are successfully recapitulated by the spatial model. Quantitative analysis of spatial SP model, simulating basal clonal growth in mouse oesophagus (*r* = 0.1, *ρ* = 0.65, *λ* = 1.9/week, from [2]). a: Progenitor cell population remains largely constant. b: The proportion of clones decreases over time, following 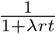. c: The average clone size increases over time, with slope *τ* = *rλ*/*ρ*. d: Clone size distribution scales over time. e: The clone size distribution adopts a broader shape over time. Data correspond to mean values across 100 simulations. Shaded areas correspond to SD.

### Advantageous mutations show logistic-like growth due to competition and feedbacks

Having confirmed that neutral systems continue to behave as expected, we moved onto examining the growth of advantageous mutations. *p53* mutations are found to be enriched in tumours, and in mouse back skin, induced *p53* mutant clones show super-linear growth and partial takeover of the tissue after 12 months [16]. This is enabled by an imbalance (*δ*) in the single progenitor model where divisions resulting in pairs of progenitor cells become more likely than divisions giving differentiated daughters that leads to exponential growth of clones. Surprisingly however, tissue takeover slows over time suggesting that other factors influence the clone takeover in a wild-type tissue. In contrast, *DN_Maml1* expression confers an advantage to mutant cells that leads to complete and rapid tissue takeover, before the tissue returning to a new homeostatic state.

Initial simulations of the mutant *p53* colonisation of the epidermis modified a set of wild-type cells to have an imbalance in symmetric cell fates. A range of levels of mutant induction was tested, reflecting the relative uncertainty in calculating the initial induction level from the exponential growth relative to the wild type situation. Without further additions or fitting of the model, *p53* clone growth and tissue takeover matched experimental observations, showing an initial exponential growth slowing over time (Fig 3a,b). Visualisation of the tissue showed increased packing of the mutant cells and an increased abundance of progenitor cells, which over time lead effectively to the growth becoming logistic. To confirm that the tissue size was acting as a limiting factor and not the hexagonal grid, the model was re-implemented as a rule based model with a carrying capacity. These simulations confirmed that whilst the presence of the grid slowed growth slightly, the growth curves were ultimately logistic-like (Fig 3c).

**Fig 3.**
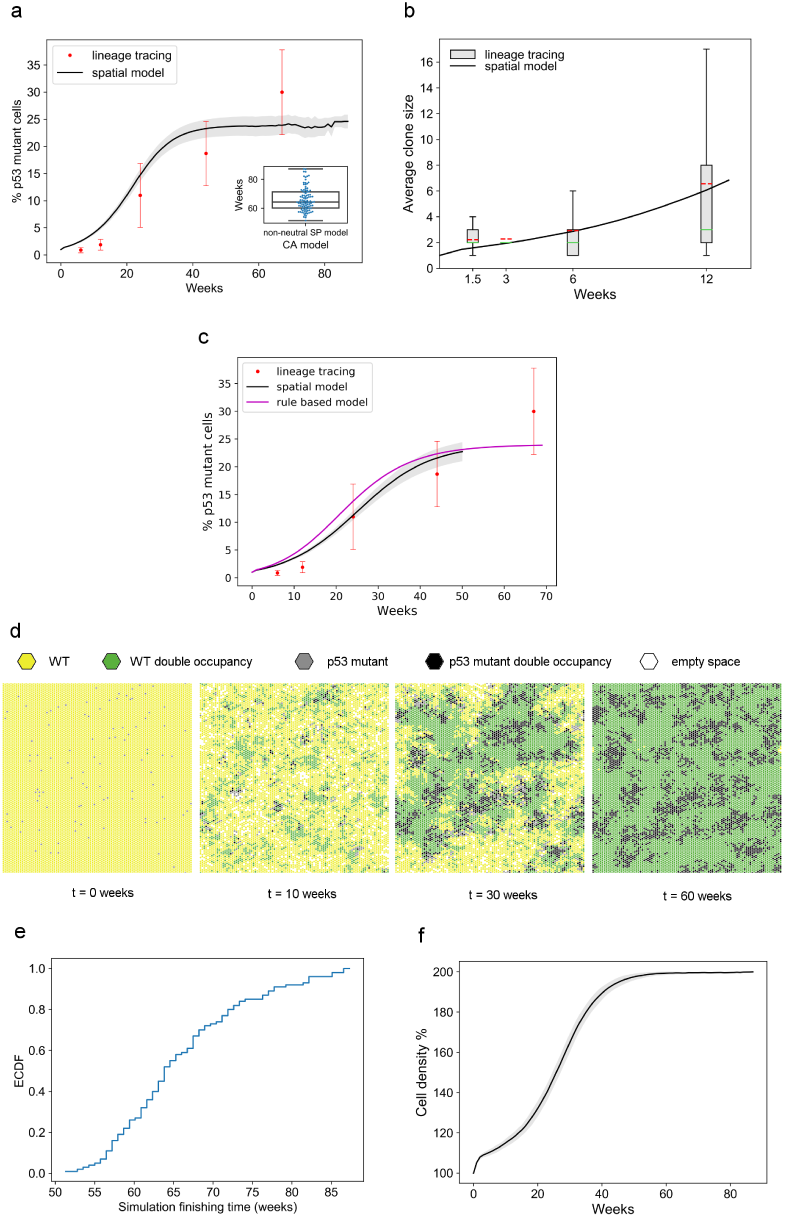
Spatial simulations of *p53* mutant growth. Quantitative analysis of the spatial non-neutral SP model including fate bias, simulating *p53* mutant growth in the basal layer of back epidermis (*r* = 0.06, *ρ* = 0.77, *λ* = 1.16, *δ* = 0.95, from [16]). a: Simulated and observed percentage of *p53* mutant cells, inset: last simulated timepoint of each repetition (100 simulation repetitions were performed). b: The simulated *p53* average clone size matches observations, box plots correspond to experimental clone counts, dashed red lines: mean, solid green line: median. c: Percentage of the *p53* mutant population in spatial and rule based simulations. The two model types display similar behaviour. Rule based simulations were performed using Bio-PEPA, a framework for modelling biochemical networks [28]. d: Typical simulation time lapse of the SP spatial model of *p53* mutant growth. Mutant behaviour fills grid and slows growth across the basal layer, leading to 100% increase in cell density. e: Empirical cumulative density function (ECDF) of last simulated timepoint across 100 repetitions. f: Cell density increases up to 100%, A,B,D,F: Data correspond to mean values across 100 simulations. Shaded areas correspond to SD.

Despite matching experimental data well, the simulations presented a new problem. As the population of progenitor cells and double cells grows over time, the overall tissue turnover slows as cells no longer have space to divide (Fig 3d). This is particularly notable in the mutated regions which become packed earliest, and is not well supported by biological evidence which suggests that the tissue is otherwise morphologically normal. This has practical implications for the model; simulations can finish within the lifetime of the mouse when the tissue becomes packed with double occupancies (Fig 3e), whereas the tissue is found to have an increase in cell density of only ~ 10% (Fig 3f).

To address these issues, we extended the model to explicitly enable feedbacks that limit growth on the basis of crowding. Cell density increases in large areas replaced by mutant cells, and is coincident with a return to neutrality in experiments [9]. Two classes of feedback were tested based on the subpopulation of cells that respond to crowding. In one class all cells respond to crowding by promoting differentation (Fig 4a). In the second class only mutant cells respond to crowding by reducing their propensity to symmetric division giving dividing cells (Fig 4b). If all cells respond to crowding, it implies that the genes responsible for coordinating this response either act independently to or upstream of the mutated gene, that are active in the whole population. Alternatively, if only mutated cells respond to crowding, it implies that the genes responsible for coordinating the response act on targets of the mutated gene that are not active in the wild-type subpopulation. We therefore term these mechanisms “upstream” and “downstream” responses to crowding respectively to reflect the implied relationship between the crowding responders and the mutated gene.

**Fig 4.**
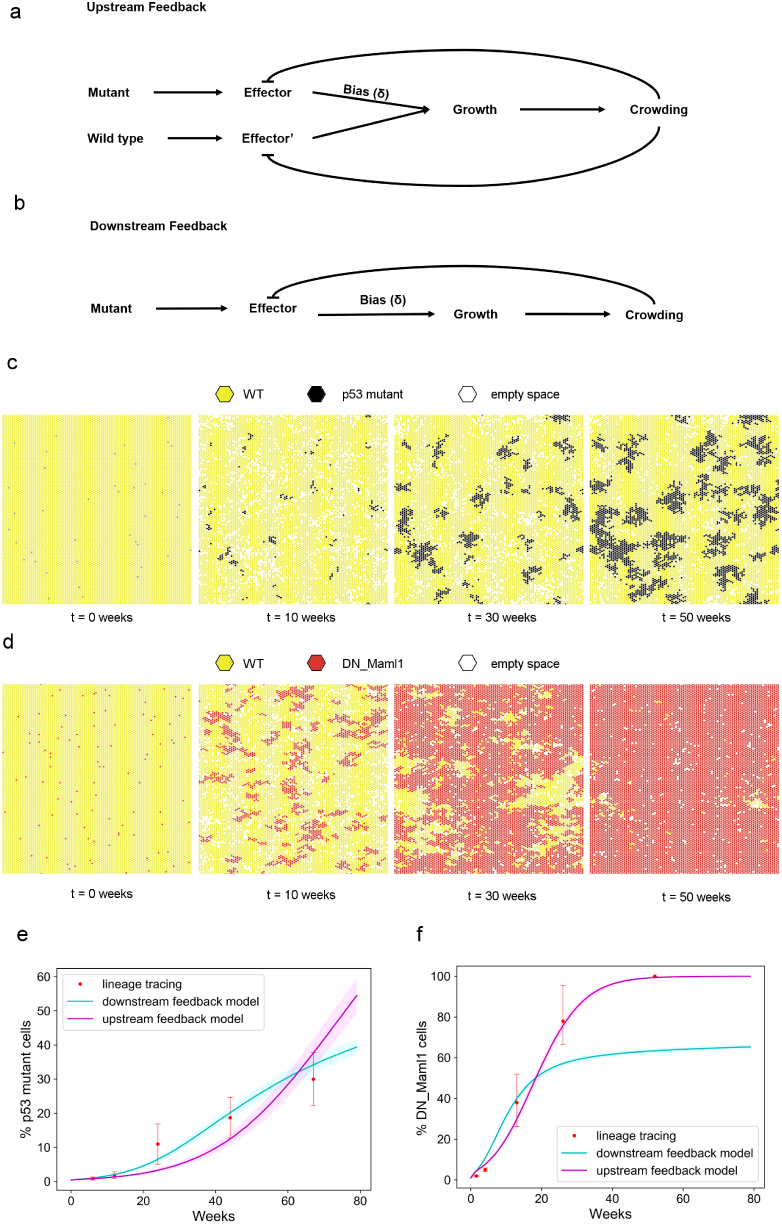
*p53* and *DN_Maml1* growth is explained by two distinct feedback mechanisms. The “upstream” (a) and “downstream” (b) feedback models simulating c: *p53* mutant growth in back epidermis (SP parameters taken from [16]) and d: *DN_Maml1* growth in oesophagus (SP parameters taken from [9]). *p53* mutant growth dynamics is appropriately recapitulated by the “downstream” feedback model whereas *DN_Maml1* by the “upstream” feedback model. e,f: Tissue takeover plots show the simulated and observed percentage of mutant cells. Data correspond to mean values across 100 simulations. Shaded areas correspond to SD. Simulation time lapses show *p53* mutant growth with the “downstream” feedback (crowding threshold was set to 7 cells) and *Maml1* growth with the “upstream” feedback (crowding threshold was set to 6 cells, *δ′* = 1).

Both approaches were tested at a range of crowding thresholds and induction levels. The upstream model enhanced tissue takeover, and no longer matched tissue takeover by the end of the mouse lifetime. In contrast, the downstream model followed the growth curves correctly and was broadly robust to the specific threshold selected (Fig 4c,e). Furthermore, simulations no longer finished in the mouse lifetime and tissue turnover remained roughly constant suggesting that the growth advantage of the mutant *p53* cells is sensitive to crowding, effectively acting as a negative feedback to the mutant cell population.

Having shown that the *p53* mutant cell growth can be described by a simple “downstream” feedback, we revisited and retested *DN_Maml1* transgenic mutations in mouse oesophagus. Testing with both feedbacks, we found that the “upstream” model was required to capture the rapid takeover. This is consistent with prior observations that showed through EdU labelling experiments that wild type cells neighbouring mutant *DN_Maml1* clones were induced to differentiate and stratify [9]. Visual examination of the underlying simulations suggested that the difference in growth patterns between the upstream and downstream models arises as in the upstream model the reduction in *δ* in all cells had the effect on ensuring that cells continued dividing, but maintained the relative advantage of the mutant clone (Fig 4d,f). This effect, where *DN_Maml1* mutation gives a constant fitness advantage whilst the *p53* mutant advantage is transient, was subsequently confirmed using a spatial Moran model. Here each cell has a pre-defined fitness, and at each step of the simulation a cell has an opportunity to replace its neighbour based on their relative fitnesses. We find that whilst *DN_Maml1* mutations can be accurately modelled using the Moran model, *p53* clone growth could not be accurately modelled with a single fitness (S2 Fig, S3 Fig).

## Competition

Having shown that both sets of mutations can be understood in terms of the interactions of the underlying pathways and the tissue, we sought to assess how a single model could enable both feedbacks behaved and how the mutations coexist and interact in the tissue. In humans the prevalence of mutations varies between normal oesophagus and the cancers that derive from it. *p53* mutants are present the great majority of cancers but are typically present in less than 10% of normal cells [12, 13]. This implies cancers develop from the *p53* mutant population. In contrast *NOTCH1* mutations are several fold more common in normal oesophagus than in cancers, hinting that wild type *NOTCH1* may promote malignant transformation. To explore the interactions of the mutations we designed a series of different mutational events to consider how the different mutations interact; co-induction, timed inductions, and variable levels of relative induction. As *DN_Maml1* represents a gene with control over several Notch related genes, we modelled the effect of *NOTCH1* mutations as an increase in the imbalance combined with the measured increase in division rate, in order to examine how the different growth behaviours in the tissue alter the competition.

We found that in all scenarios *DN_Maml1* mutant clones eventually took over the tissue, excluding *p53* clones from the tissue (Fig 5a,b). *p53* clones that had grown for extended periods before *DN_Maml1* mutations were introduced regressed more slowly, as the larger clones were more slowly outcompeted but ultimately the constant fitness advantage offered by *DN_Maml1* mutations eventually led to the loss of *p53* clones. However, a different *p53* favouring competition scenario, where *p53* mutants were introduced at a substantially higher proportion, led to a higher rate of *p53* clone regression compared to scenarios of delayed *DN_Maml1* induction (Fig 5c).

**Fig 5.**
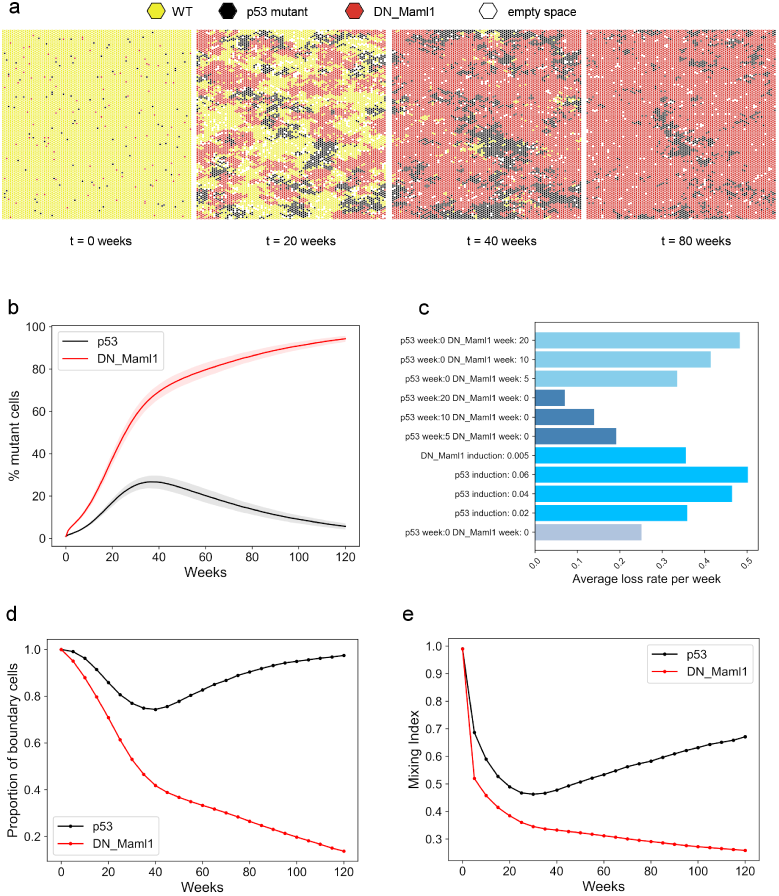
*DN_Maml1* mutations outcompete *p53* mutant cells in competition simulations of oesophagus. a: Typical competition simulation time lapse assuming mutant co-induction and same induction level. b: Tissue takeover percentage of *p53* and *DN_Maml1* mutations, when both are introduced in the tissue. c: Average percentage of *p53* mutant takeover loss rate across all simulated competition scenarios. Calculations measured the difference in tissue takeover between the week when takeoverwas the highest until the final week of the simulation. d: Proportion of boundary cells in *p53* and *DN_Maml1* mutant cells. e: Proportion of neighbours of a different mutation type or wild-type (cell mixing index) as computed for *p53* and *DN_Maml1* mutations. The cell mixing index was calculated for each mutant cell and the values were averaged over each mutation. Data correspond to mean values across 100 simulations. Shaded areas correspond to SD. Parameters used: *DN_Maml1*: SP parameters taken from [9], *δ* = 1 (from [9]), crowding threshold: 6 cells, *δ′* = 1, *p53*: SP parameters taken from [2], *δ* = 0.95 (from [16]), crowding threshold: 6 cells.

In seeking to quantitatively describe potential distinct spatial properties of the two competing mutant populations, we observed that the majority of *p53* mutant cells were consistently found in boundaries with *DN_Maml1* or wild-type populations, implying that *p53* mutant cells were not able to form coherent groups in space. On the contrary, the proportion of boundary *DN_Maml1* mutant cells progressively decreased and was almost minimized at later time points, as they colonized the entire grid (Fig 5d).

Furthermore, the proportion of *p53* mutants’ neighbours belonging to a different type (cell mixing index) was consistently higher, indicating that *p53* mutant cells were more likely to share junctions with different cell types and therefore tended to be more dispersed (Fig 5e). This is consistent with experimental observations [16].

The quantified distinct spatial features might be able to explain the behaviour observed across different competition scenarios. When *p53* mutants are introduced in the grid in much higher numbers compared to their *DN_Maml1* competitors, they would expand and form aggregates more easily and faster. However, as they cluster in space and tend to appear in more crowded environments and encounter more *p53* mutant neighbours, they would start losing their innate proliferating advantage. The subsequent contact with *DN_Maml1* mutant cells which start expanding substantially in the following time points would lead to their loss as *DN_Maml1* mutant population is more aggressive.

## Discussion

The growth and competition of cells in epithelial tissues plays an important role in both tissue homeostasis and the robustness to pre-cancer mutation. Epithelial tissues are environments of tightly packed cell populations which constantly compete for limited space in a perfectly coordinated manner to ensure homeostasis. The emergence of non neutral mutations may provide a competitive fitness advantage which allows the dominance of the fitter population. Spatial competition of accumulated non neutral mutations within tissues could potentially drive tumourigenesis.

Here, we explored the growth and competition of non neutral mutations in stratified squamous epithelia using spatial models. Mutations in the tumour suppressor gene *p53* and inhibition of the Notch pathway through *DN_Maml1* knock out were modelled. The studied dynamics of *p53* and *DN_Maml1* clones in mouse epithelia has been found to be highly distinct. Whilst *DN_Maml1* knock outs are able to dominate the whole tissue [9], *p53* mutants initially expand but they eventually occupy no more than 30% of the tissue [16]. We showed that in order to recapitulate these behaviours whilst maintaining tissue turnover the spatial models need to take account of feedbacks between neighbouring cells in the tissue. These are based on spatial regulatory rules between surrounding cells in order to locally balance cell density.

We demonstrated that *p53* clone growth approximates a logistic curve, but that without including limitations on mutation induced expansion the overall proliferation rate of the tissue drops due to space restrictions. In contrast, the ability of *DN_Maml1* mutations to take over the progenitor cell population reflects a feedback that affects both mutant and wild-type cells equally. Thus, for reproducing *p53* mutant behaviour, we used a model where the feedback response was applied only to mutant cells, indicating activation downstream to *p53* mutations (“downstream” feedback). Such model appears consistent with the *p53* mutants’ adaptable fate, suggested by their observed incomplete tissue takeover. On the contrary, we modelled *DN_Maml1* dynamics using a feedback mechanism affecting all cell populations (“upstream” feedback), consistent with the ability of *DN_Maml1* cells to evict their neighbours during their early growth [9]. These spatial models of *DN_Maml1* and *p53* mutant growth could provide a mechanistic explanation of the observed distinct behaviours of the two mutant types, suggesting an increased sensitivity of *p53* mutant clones to cell density.

*p53* and *DN_Maml1* competition in space was also explored. A striking effect resulting from the spatial interaction of the two mutations in a wild-type background is that *p53* mutant cell population was always outcompeted by *DN_Maml1* population and appeared to shrink. The winning behaviour of *DN_Maml1* cells was consistent across a range of competition scenarios, where the two mutant populations were introduced in the simulations at different time points or relative inductions. A further quantitative comparison of the spatial features of mutant competition revealed distinct growth patterns. Whilst *DN_Maml1* cells formed coherent aggregates in space which progressively expanded at the expense of their neighbours, *p53* mutant cells initially clustered but eventually appeared more dispersed and less cohesive, consistent with experimental observations [16]. The consistent winning behaviour of Notch mutations comes in contrast to a recent study, suggesting that the persistence either Notch or *p53* mutant clones is determined by time [24].

Taken together, these findings suggest that the distinct tissue takeover outcomes may be attributed to the differential response of *p53* and *DN_Maml1* mutations to crowding. Moreover, according to the competition spatial simulations, the sensitivity of *p53* mutant cells to increased cell density environments might be the cause of their loser status. The crowding sensitivity as a hallmark of loser cells is also supported by previous studies [7, 17, 25]. Furthermore, the increased dispersion of *p53* mutants at later time points in competition simulations, indicates that this spatial emergent property might be a result of their mixing with *DN_Maml1* populations. This would be consistent with previous studies in drosophila tissues, correlating the probability of loser cell elimination with the surface of contact shared with winners [26]. Further experimental work on *p53* and *DN_Maml1* competition in mouse epithelia would shed more light on how these two mutants interact in mammal tissues.

The two distinct feedback mechanisms that describe mutant dynamics may suggest the distribution of mutations observed in human datasets. Recent studies identified high incidence of many of the frequently found mutated genes in tumours such as *p53* and *NOTCH1* in healthy human skin and oesophagus [12–14]. Strikingly, a higher frequency of *NOTCH1* mutations is reported for healthy human oesophagus compared to oesophageal squamous cell carcinomas whilst the opposite applies to *p53* mutations. Considering the consistent winning behaviour of *NOTCH1* over *p53* mutants in our competition simulations and given the paucity of *NOTCH1* mutations in tumours, it is tempting to speculate that the aggressive fitness of *NOTCH1* may offer a tumour-protective effect [27] under some circumstances. If however the mutations may act combinatorially, *p53* mutations that precede a *NOTCH1* mutations may both expand and promote cancer development. Given that *p53* clones persist in aged samples from healthy patients, this may suggest that either the speeds of clonal expansion are slower in human tissues or that other processes, such as secondary mutation, enable their persistance. The relationship between these genes, and the specific environment therefore represent an important topic of future studies.

## Acknowledgements

We thank the Hall and Jones groups for discussions. This work has been supported by the Royal Society (URF to BAH grant no. UF130039, studentship to VK), Medical Research Council (Grant-in-Aid to the MRC Cancer unit), the Harrison Watson Fund at Clare College, Cambridge (M.W.J.H.), Cancer Research UK Programme (P.H.J., grant no. C609/A17257), & the Wellcome Trust (to the Wellcome Sanger Institute, 098051 and 206194).

## Supporting information

**S1 Fig.**
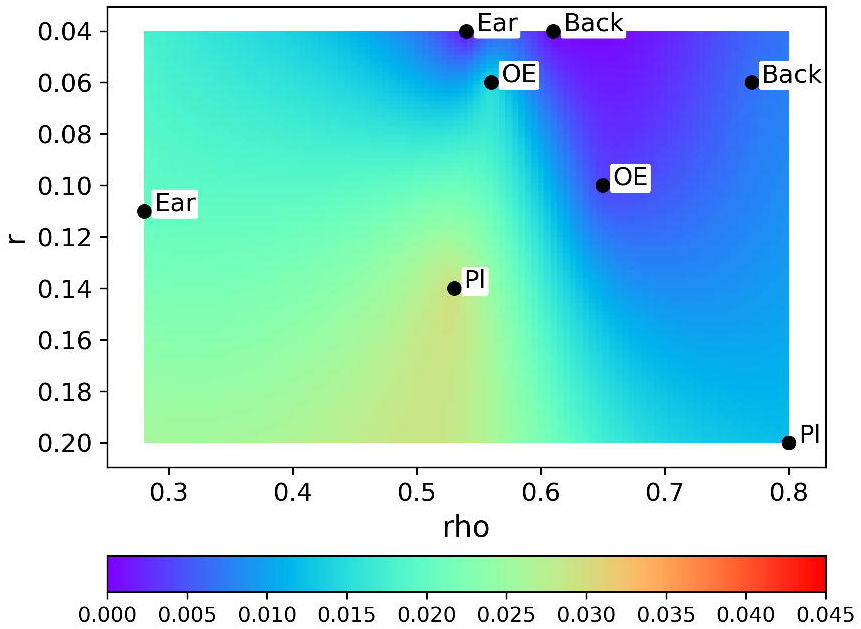
Line slope VS 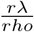. The average clone size increases linearly with slope 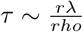 across different datasets. Figure shows the absolute difference between 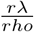 ratio for each dataset with known *r*, λ, *rho* values and the slope of the line of average clone size.

**S2 Fig.**
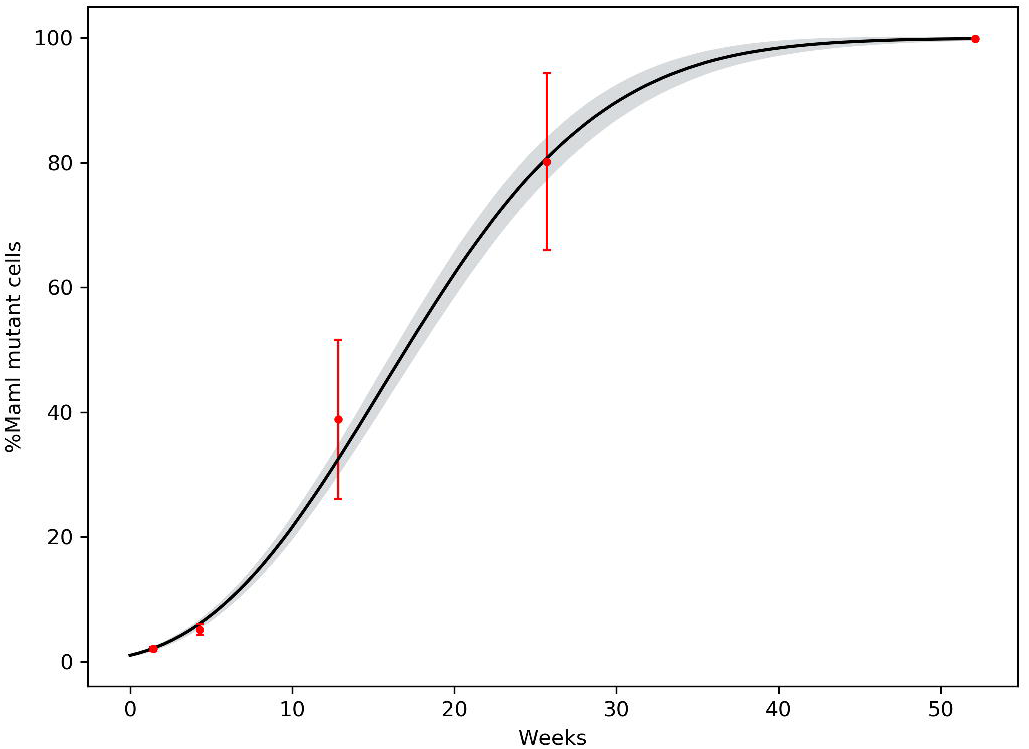
*DN_Maml1* Moran-style 2D model. The Moran process is able to reproduce tissue takeover accurately. This finding is consistent with the observation that “upstream” feedbacks maintain the relative fitness advantage of the clones.

**S3 Fig.**
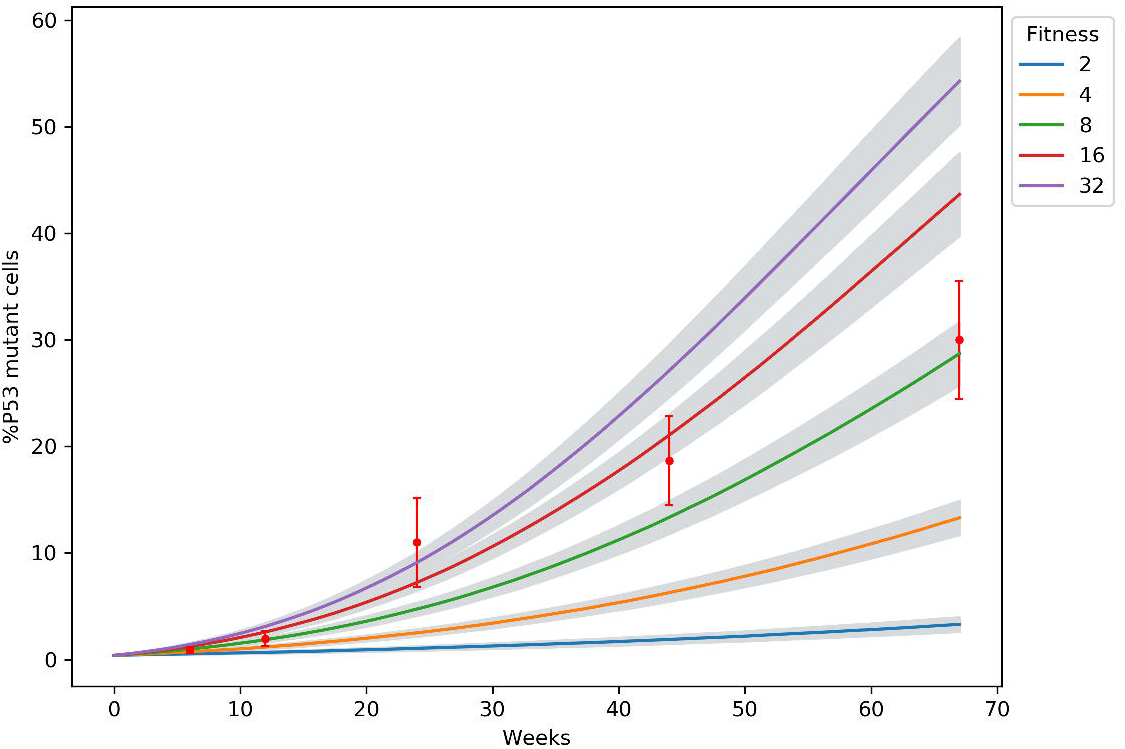
*p53* Moran-style 2D model. The Moran process is unable to reproduce tissue takeover, either overestimating or underestimating takeover depending on the specific choice of parameters. This observation is consistent with feedbacks leading to phenotypic plasticity of the *p53* clones as suggested in the “downstream” model.

